# JarrVis: Visualising Taxa-function relationships from meta-omic data

**DOI:** 10.1101/2025.04.13.648596

**Authors:** Dhwani K. Desai, André M. Comeau, Morgan G.I. Langille

## Abstract

High-throughput sequencing has necessitated and facilitated the development of various computational tools for dissecting the taxonomic structure and molecular function of microbial communities residing in diverse ecological niches. A wide variety of tools and protocols have been developed to process amplicon sequencing as well as meta-omic (metagenomic, metatranscriptomic and genomic) data to generate relative abundances of taxonomic groups or functional categories on a per-sample basis. A key output from many of these protocols is a stratified relative abundance table that specifies the taxonomic contribution to each function. However, there are a very few tools which can effectively visualize the different taxa-function relationships from these stratified outputs.

Here we introduce an R Shiny application called JarrVis (Just Another stRatified Rpkm VISualizer) which can visualize the taxa-function relationships resulting from different types of microbiome data. JarrVis can visualize relationships between samples (which can be combined based on metadata categories), the taxa that are detected in these samples (at any given taxonomic level) and the functions encoded by these taxa in an interactive interface.

We utilized JarrVis to visualize and examine taxa-function relationships in 1) a 16S amplicon time-series spanning 4 years with samples collected weekly, 2) functions associated with microbial cobalamin biosynthesis and uptake in metagenome-assembled genomes from marine metagenomes and 3) a set of metatrascriptomes from gut samples of patients with Crohn’s Disease, Ulcerative Colitis and non-IBD controls. Our analysis was able recapitulate already published taxa-function relationships as well as discover novel insights from these publicly available datasets. JarrVis and related scripts and data are available at https://github.com/dhwanidesai/JarrVis.

## Introduction

Metagenomic and metatranscriptomic (meta-omic) sequence analysis, including microbial functions inferred from amplicon sequencing (Douglas *et al*., 2020), results in stratified taxonomic and function relative abundance tables in terms of reads per kilobase per million (RPKM) or other such normalized values. Our in-house bioinformatics workflow (Microbiome Helper) for bulk metagenomic analysis also produces RPKM values of microbial functions stratified by contributions from taxonomic groups (Comeau *et al*., 2017). Several tools exist for visualization of the taxonomic or functional diversity in meta-omic samples (Meyer *et al*., 2008; Ondov *et al*., 2011; Huson *et al*., 2007; Foster *et al*., 2017; Caspi *et al*., 2020; Mcmurdie and Holmes, 2013). These tools enable an exploration of unstratified taxa or function relative abundance tables individually to identify taxonomic or functional signatures in groups of samples.

However, there is a distinct lack of availability of software for visualizing such taxa-function relationships in the context of the treatment/control groups or environmental niche groups (Sudarikov *et al*., 2017; McNally *et al*., 2018; Murat *et al*., 2021; Bik and Inc., 2014; Peeters *et al*., 2021). Here, we introduce JarrVis, an interactive R shiny app (Chang *et al*., 2021; R Core Team, 2021), which provides a visual exploration of the processed metagenomic, metatranscriptomic or genomic data in terms of taxa-function relationships and how they relate to specific environmental niches (groups of samples). We demonstrate its utility to uncover novel taxa-function signatures corresponding to specific environments from published meta-omic datasets.

## Methods and Features

Ease of use and reproducibility: JarrVis is written in R Shiny and can be run in RStudio either as a standalone app or from its GitHub Gist available at https://gist.github.com/dhwanidesai/943ff5fdbd94815cc27f302d9f56ff0b. This ensures reproducibility as there is no need to individually install any dependencies.

Datatype agnostic: JarrVis can display Sankey diagrams of taxa-function relationships from a wide variety of data types including stratified metagenomic or metatanscriptomic RPKM tables, PICRUSt2 predicted functions from amplicon data or even genomes (complete or assembled from metagenomes). Any data that has the format of relative abundances of functions stratified by taxa and grouped according to certain criteria (sample metadata, genomes isolated from specific niches etc.) can be visualized in JarrVis (Figure 1A).

**Figure 1.**
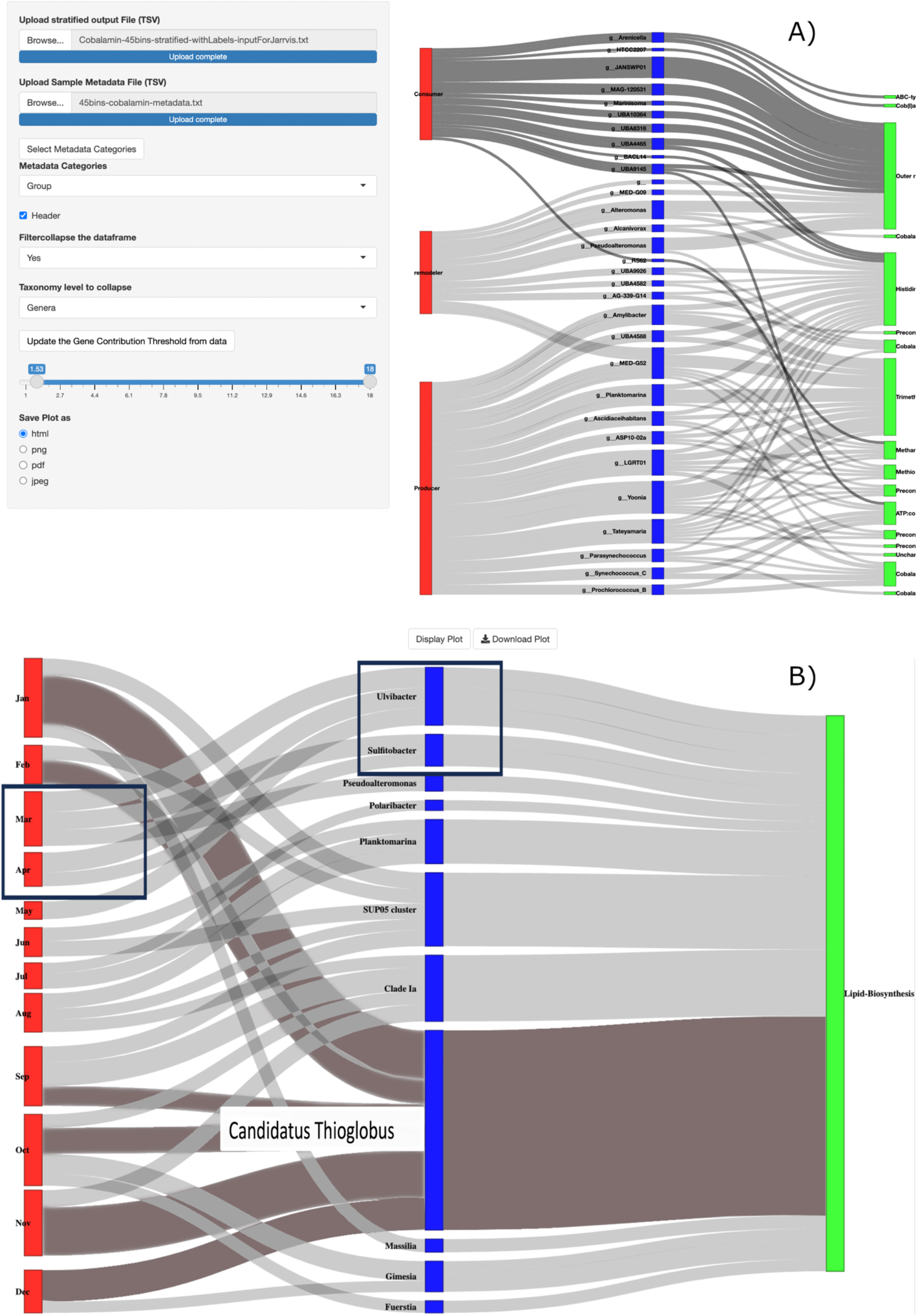
A) The user interface for JarrVis depicting the input controls and the output Sankey diagram. B) JarrVis output showing the month-wise shift in microorganisms having potential for lipid biosynthesis in the weekly time-series samples from Bedford Basin, for the year 2014, collected at a depth of 60 m. The grouped sample nodes are shown in red, the taxonomic nodes in blue and the functional nodes are depicted in green.

Interactive Sankey diagrams: The displayed Sankey plots can be interactively manipulated. For example, right-clicking on the sample group nodes can be used to highlight the linkages all the way through to the taxonomy nodes and the corresponding function nodes. Right-clicking on the function nodes can highlight the linkages all the way through to taxonomy nodes and the corresponding sample group nodes. This makes it easier to highlight linkages between specific functions and the corresponding sample groups and vice-versa. You can left-click and drag the nodes to change their positions. Hovering over nodes or links displays the underlying RPKM values.

Input: The main input file format for JarrVis is a tab delimited file with four columns corresponding to sample ID, complete taxon name with 6 levels of taxonomy, gene name or function name and the normalized contribution value (for example RPKM), signifying the relative abundance of the given gene or function stratified by the taxonomic categories. In addition, a tab delimited metadata file specifying how the samples should be grouped is also required. The JarrVis interface provides options to select and filter the stratified input file interactively by searching for specific taxonomic categories or functions. Similarly, the metadata category to group the samples can also be interactively selected. Once the dataset is filtered and samples grouped by metadata category, the RPKM range can be re-adjusted using the minimum and maximum values from the data, and subsequently, interactively changed to specify a range of RPKMs to display (for example to filter out low-abundance “noise”).

Output: The generated Sankey plots can be downloaded in HTML, PDF, PNG or JPEG formats. The HTML output can be interactively manipulated in a browser to highlight important linkages and to re-arrange the sample, taxa or function nodes and the resulting manipulated plot exported as a PDF file.

### Seasonal change in microbial taxa associated with Lipid Biosynthesis in the Bedford Basin

In a previous study, 16S amplicon sequences were analyzed for the years 2014 to 2018 as part of a weekly time-series in the Bedford Basin (Raes *et al*., 2022). Potential functions were predicted from 16S amplicon sequence variants (ASVs) using PICRUSt2.

Statistical analysis of predicted functions revealed lipid biosynthesis as having significant positive correlation with spring blooms (Raes *et al*., 2022). We used JarrVis to reanalyze the stratified output linking the taxonomic and functional profiles of relative abundances for 52 weekly samples for the year 2014 collected at a depth of 60 m. The stratified relative abundance was filtered to include only the function lipid biosynthesis and the samples were grouped by month to observe the seasonal effect. A visual exploration in JarrVis showed that distinct microbial taxa were major contributors to the lipid biosynthesis function at different time points during the year. For example, Candidatus *Thioglobus* (highlighted in grey), was the major contributor during the late fall and winter months from September to February, whereas spring contribution (March and April) came mainly from *Ulvibacter* and *Sulfitobacter* spp. (Figure 1B).

In another case study, we re-analyzed stratified taxonomic and functional profiles from gut microbiome samples of patients with Crohn’s Disease (CD) or Inflammatory Bowel Disease (IBD) as well as healthy subjects without any disease (Schirmer *et al*., 2017). These profiles were output by HUMAnN2 (Franzosa *et al*., 2018). We focused on 28 metagenomes from 6 individuals (2 each for IBD, CD and non-IBD) collected at approximately 0, 100, 200 and 300 days. Visualization with JarrVis re-captured certain microbial signatures associated with Crohn’s Disease and Ulcerative Colitis, for example, higher relative abundance of *B. vulgatus* and *F. prausnitzii* in the two disease groups (Supplementary Figure 1). We also observed that the galacturonate degradation pathway (GALACTUROCAT PWY) was almost solely associated with *F. prausnitzii* for the CD samples (Schirmer *et al*., 2017). One interesting observation was the presence of micronutrient pantothenate and coenzymeA (CoA) biosynthesis-related pathways in the genus *Alistipes*, associated with non-IBD healthy samples (Supplementary Figure 1).

IBD fecal microbiomes have been correlated with reduced dietary intake of pantothenate, a precursor of CoA, which in turn, is required for the production of beneficial short-chain fatty acids (SCFAs) (Kundra *et al*., 2021; Weng *et al*., 2019; lloyd-Price *et al*.; Franzosa *et al*., 2019; Santoru *et al*., 2017; Louis and Flint).

This and other case studies demonstrating the utility of JarrVis are described in detail on the JarrVis GitHub wiki (https://github.com/dhwanidesai/JarrVis/wiki).

## Conclusion

We present here an R Shiny app for interactive visualization of taxa-function relationships derived from meta-omic or comparative genomic data. The tool JarrVis and the accompanying scripts, in addition to the relevant documentation, is available from https://github.com/dhwanidesai/JarrVis.

