## Supplementary Figure 1 for "JarrVis: Visualising Taxa-function relationships from meta-omic data"

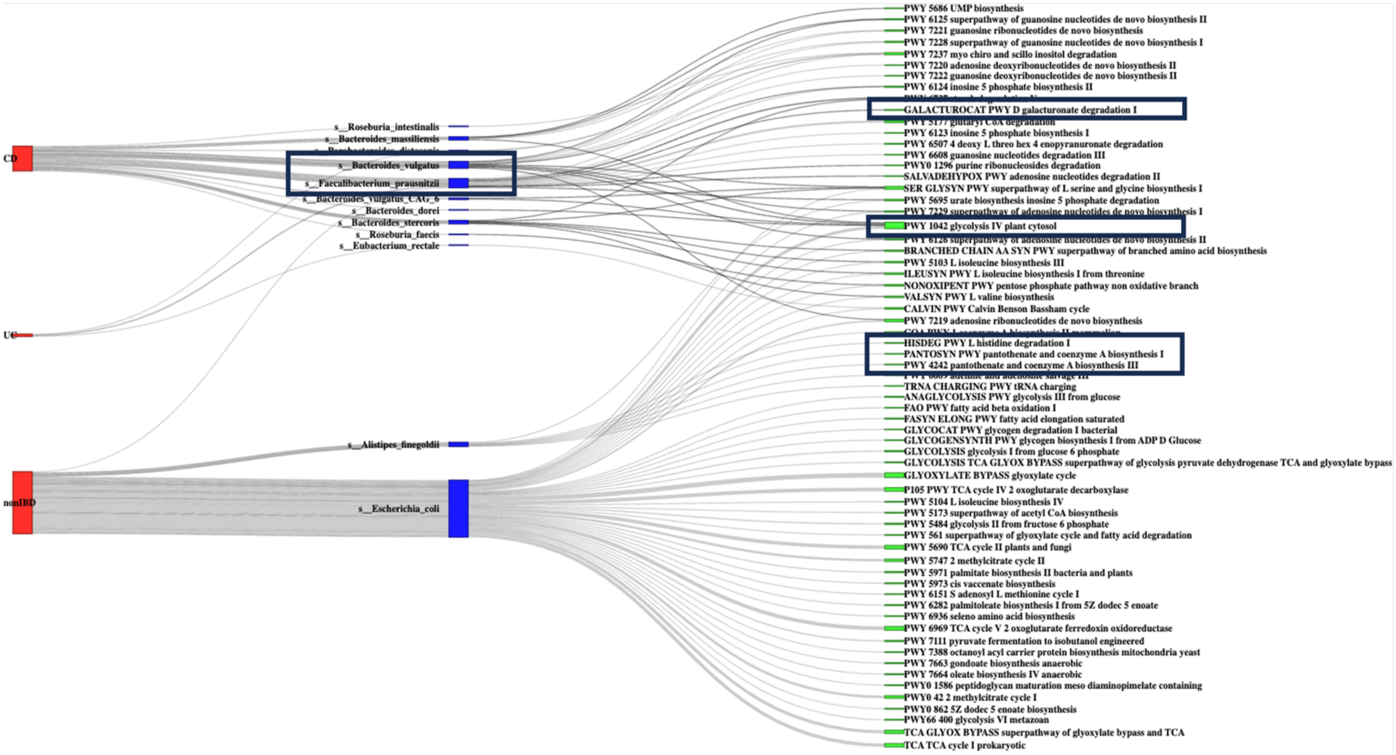


Supplementary Figure 1: JarrVis output showing the microbial diversity linked to pathways in gut metagenomic samples from patients with Crohn’s Disease (CD), Inflammatory Bowel Disease (IBD) and healthy controls (nonIBD).
